# Visible neural networks for multi-omics integration: a critical review

**DOI:** 10.1101/2024.12.09.627465

**Authors:** David Antony Selby, Rashika Jakhmola, Maximilian Sprang, Gerrit Großmann, Hind Raki, Niloofar Maani, Daria Pavliuk, Jan Ewald, Sebastian Vollmer

**Author notes:** These authors contributed equally to this work (David Antony Selby, Rashika Jakhmola, Maximilian Sprang).

## Abstract

Biomarker discovery and drug response prediction is central to personalized medicine, driving demand for predictive models that also offer biological insights. Biologically informed neural networks (BINNs), also known as visible neural networks (VNNs), have recently emerged as a solution to this goal. BINNs or VNNs are neural networks whose inter-layer connections are constrained based on prior knowledge from gene ontologies pathway databases. These sparse models enhance interpretability by embedding prior knowledge into their architecture, ideally reducing the space of learnable functions to those that are biologically meaningful. In this systematic review—the first of its kind— we identify 86 recent papers implementing such models and highlight key trends in architectural design decisions, data sources and methods for evaluation. Growth in popularity of the approach is apparently mitigated by a lack of standardized terminology, tools and benchmarks.

## 1 Introduction

High-throughput technologies have transformed biological research, enabling the collection of large-scale data from different molecular layers and leading to the emergence of multi-omics, an approach that combines information from diverse sources, such as genomics, transcriptomics, proteomics and metabolomics [1]. This allows researchers to analyse complex interactions and regulatory mechanisms that drive cellular function, disease progression and response to therapeutic interventions, with the potential to identify novel biomarkers and better understand how genetic and environmental factors influence disease [2].

However, multi-omics analysis is fraught with challenges due to high dimensional, heterogeneous data, requiring methods capable of modelling non-linear relationships across multiple processes. Modern machine learning (ML) techniques, such as deep learning, offer superior predictive capabilities at the expense of interpretability. In this context, even explainable AI (XAI) metrics may lack clear or robust biological interpretations [3–5].

To meet this demand, visible neural networks (VNNs), also known as biologically-informed neural networks (Binns), have gained prominence. Unlike conventional neural networks (NNs), which learn relatively unconstrained functional approximations, VNNs incorporate prior knowledge directly into their architecture: previously ‘hidden’ nodes map directly to entities such as genes or pathways, with inter-layer connections constrained by their ontology (see Figure 1).

**Figure 1.**
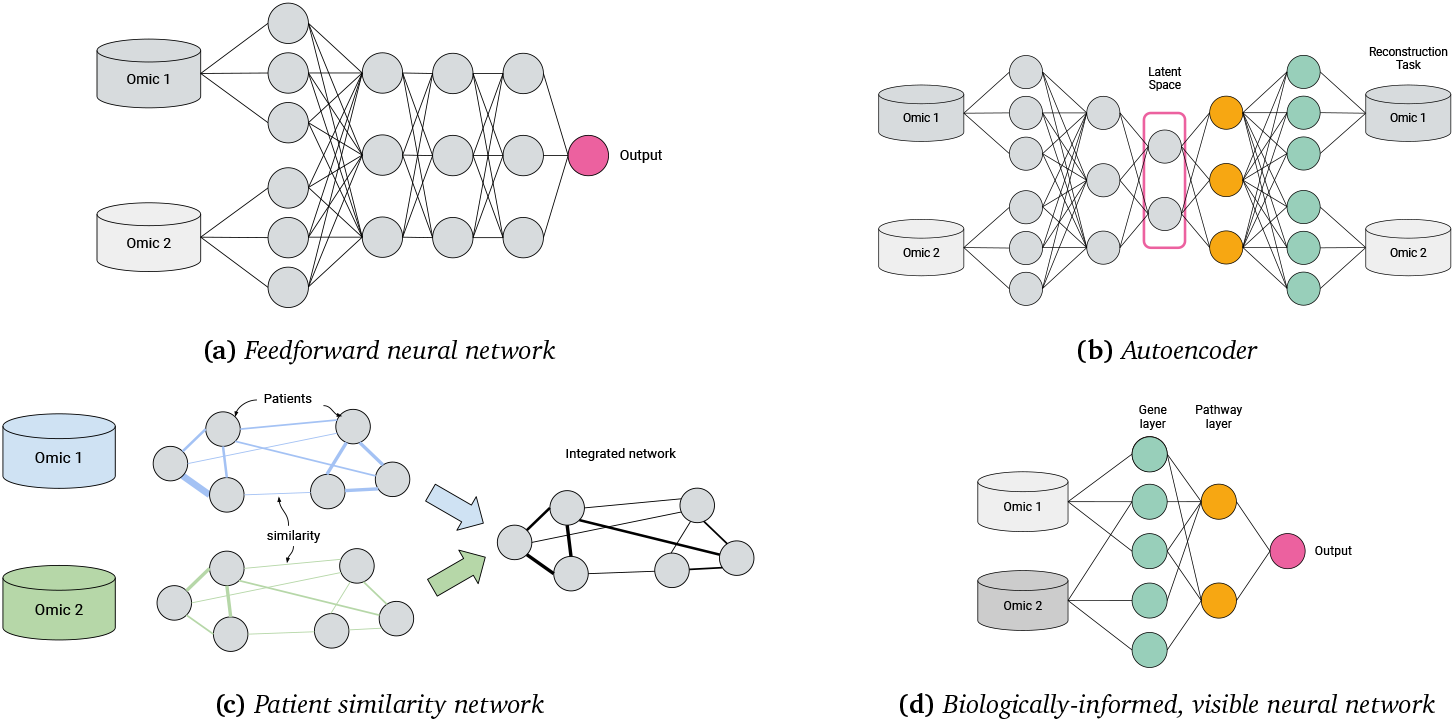
Machine learning approaches to multi-omics integration

VNNs have the potential to enhance biomarker discovery and drug development by learning relationships between genes, pathways, drugs and disease phenotypes. Various VNN models have been proposed, but standard terminology and clear design practices—fostering scientific reproducibility, robustness and generalizability—do not yet exist.

Several existing survey papers offer a high-level overview of interpretable deep learning in omics, but without focussing on detailed comparisons of biologically-informed architectures. This review *focusses specifically on VNNs*, filling an important gap in the literature. We examine design considerations in constructing different VNN architectures and applying them to different tasks and data sources.

Our contributions are as follows: a taxonomy of Binn architectures, a critical appraisal of the dependencies, assumptions, data sources and tools involved in building Binns/VNNs; and identification of several research gaps and future work. We consider three underlying research questions:

1. How do predictions of Binns compare with non-biologically-informed models?
2. Are biological interpretations from VNNs robust to architectural design decisions?
3. Can VNNs uncover new scientific knowledge?

## 2 Background

A living cell is a complex system of interacting molecules, where metabolites and energy are used to form biomass. The cell’s regulation is guided by the central dogma of biology: DNA encodes RNA, which, in turn, encodes proteins [6]. RNA and proteins both play roles in metabolism, structure, replication and other cellular processes, interacting with each other and with DNA. These interactions can regulate cellular functions, for example as recruiting transcription factors or opening chromatin in eukaryotes [7]. Gene interactions have been extensively studied, expanding our understanding of cellular interaction networks, or *pathways* [8]. Pathways connect entities such as genes, proteins and metabolites with other tissue components, reflecting cellular functions from simple growth to complex immune response. Databases describing such molecular interactions are publicly available (see Figure 6 and section 5) representing decades of biomedical research into causal relationships.

Multi-omics analysis uses data from heterogeneous sources to describe biological processes [9, 10]. Modelling such sensitive and expensively collected data necessitates modern statistical methods that exploit existing domain knowledge to solve the ‘small data’ problem [11]. Biomarker discovery in single-omics data often relies on regularized linear models fit to a target of interest. Biomarkers are derived from signals of significant features, with enrichment analysis providing insights into their biological functions, or the features with largest coefficients identified as targets for validation in the laboratory [12].

Explainable AI (XAI) is a growing field of research that aims to produce methods to explain the reasoning behind ML models’ predictions. *Ante-hoc* explainability refers to models that are intrinsically interpretable, whilst *post-hoc* methods are a way of gaining insights about ‘black box’ model predictions in terms of their inputs [13]. Statistical models [or ‘data models’; 14], such as generalized linear models, are preferred when the main goal is inference: understanding relationships between variables that describe a natural process. On the other hand, when accurate prediction is a priority, especially when datasets are large and high-dimensional, black-box models (‘algorithmic modelling’), are preferred, and capable of learning signals from data with intrinsic structure, such as images and sequences. However, larger samples and computational resources are necessary for such methods to outperform simpler models on tabular data [15].

To simplify the learning process, the search space can be constrained through provision of additional information from prior domain knowledge. Many taxonomies exist for these methods [16–18]. Data augmentation involves generating additional data through preprocessing, e.g. varying contrast of an input image. Similarly, Yang *et al*. [19] consider metabolite perturbations as input, but enrich this data through simulation. Another method involves crafting specialized loss functions for specific domains. For instance, Jia *et al*. [20] uses a loss based on physical laws of energy distribution. Most relevant for this review is constraining the computational graph, creating an inductive bias in the neural network. Convolutional layers, for example, consider the dependence of neighbouring pixels in an image, while neural ordinary differential equations [21] account for the structure of physical processes. VNNs also fall into this category and a taxonomy is given in section 5.

Knowledge graphs (KGs) [22] can integrate data from various sources, including scientific literature. Increasingly used in bioinformatics to encode and retrieve complex knowledge, KGs describe entities— including genes, proteins, pathways, diseases, and functional annotations—across multiple levels of organization [23]. They are often built upon pathway databases, such as Reactome [24], Gene Ontology [25] or Kegg [26], that provide curated ontologies focussing on cellular processes, metabolism, interactions or signalling. These databases or networks can themselves be seen as ‘simple’ KGs with restricted set of entities and relations. Biological KGs combined with network analysis methods offer the potential to discover or explain new relationships between drugs, genotypes and phenotypes [27] and their embeddings have been employed in biomarker discovery [28], multi-omics integration [29] and drug-target discovery [30]; for example, Glue is a KG-based architecture that integrates omics features based on regulatory interactions [31].

Constructing a NN based on such databases is a form of knowledge-intensive ML. Integrating the most general KGs may result in very large NNs that fail to learn relevant patterns. Prior knowledge integration is a universality–generalizability tradeoff: compared to a dense neural network, a VNN’s function space is reduced, losing universality but generalizing better to the real-world context of the specific biological systems of interest [32]. In most cases, this is achieved by leveraging pathway databases such as Gene Ontology, Kegg or Reactome to inform the design of hidden layers in the network, ensuring that (some or all of) the model’s internal representations align with known entities and relationships. VNNs aim to emulate signalling processes, enhancing interpretability and biological relevance, thereby accurately modelling complex systems and facilitating discovery of novel insights. As Hanczar *et al*. [33] observe, this approach aims to bridge the gap between data-driven ML models and mechanistic understanding, making it particularly valuable in fields like genomics and systems biology, where omics data lack the structure of imaging or text data commonly exploited by popular deep learning frameworks.

### What’s in a name?

‘Visible neural network’ (VNN) is one of myriad terms to describe a NN model whose hidden layers and connections are constrained by a pre-specified ontology; terminology is far from standardized, making literature search difficult. ‘Visible’ emphasizes interpretability without restricting to biological applications: similarly, knowledge-primed [34], knowledge-guided [35, 36] or knowledge-based neural networks [37] or ontology-based autoencoders [38, 39] allow for hierarchical structures beyond biology. Hartman *et al*. [40], Elmarakeby *et al*. [41] and Voigt *et al*. [42] prefer ‘biologically informed neural networks’ (Binns), but this term is also used more generally [43] to describe NNs with *ante-hoc, ad-hoc* or *post-hoc* interpretability. ‘Binns’ can also refer to biologically-informed neural-symbolic methods [44–46], or biologically-inspired NNs found in connectomics [47] or neuromorphic computing [48].

## 3 Related work

VNNs are a growing sub-field at the intersection of deep learning-based multi-omics integration, interpretable ML methods for biology, and XAI more generally. Across these areas, various survey papers have been published that mention the concept of VNNs and Binns, however, most do not discuss this area in detail. An overview of these review articles, including their scope and limitations, is given in Table 1.

**Table 1.**
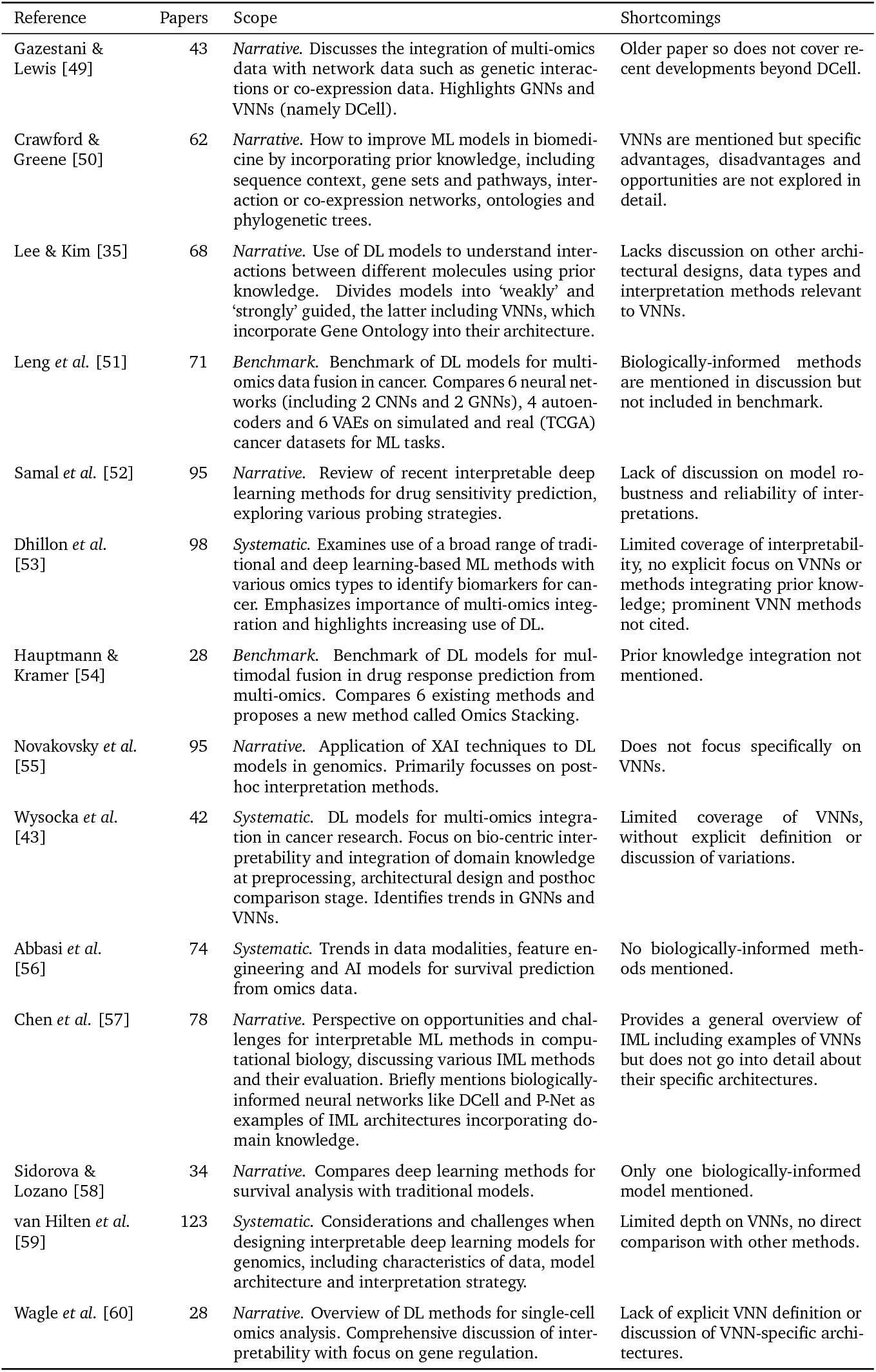
Existing review papers on explainable deep learning for multi-omics.

We consider three categories of reviews: narrative reviews, systematic reviews and quantitative benchmarks.

### Narrative reviews

Gazestani & Lewis [49] presented an early review of methods of ontology integration, including methods with hierarchical layers defined by gene ontology (i.e. VNNs), as well as network propagation models, a special case of GNNs. Crawford & Greene [50] reviewed approaches for embedding biological structures into machine learning models, including sequence encodings, network embeddings and models constrained by ontological structures. Our review specifically narrows down to VNNs that use pathways and ontologies for structured input integration. Other relevant works include Novakovsky *et al*. [55] on explainable AI in genetics, and Lee & Kim [35], which distinguishes between ‘weak’ (e.g. data-driven GNNs) and ‘strong’ biological guidance (e.g. structured VNNs). Wagle *et al*. [60] provided a review of interpretable deep learning models specifically for single-cell omics, highlighting the importance of model transparency in this context. Chen *et al*. [57] provided a broad perspective on interpretable ML in computational biology. They mention biologically informed methods and categorize them, together with attention based algorithms, as ‘by-design’ methods, which they see as naturally interpretable. Samal *et al*. [52] reviewed interpretable deep learning models for drug sensitivity prediction, exploring various ‘probing’ strategies for examining how input data is processed within the network, including some biologically-informed approaches. Sidorova & Lozano [58] asked whether deep learning methods for survival analysis outperformed traditional models, mentioning biologically-informed architectures [61, Cox-PASNet, 62, PathExpSurv].

### Systematic reviews

Wysocka *et al*. [43] conducted a systematic review of Binns. They derived a taxonomy of pre-processing, in-processing and *post-hoc* biological interpretability. Their review spans 42 publications up to 2022, sourced from PubMed and the *Web of Science*. Focussing on oncology, it covers a range of interpretability methods, including GNNs and *post-hoc* explainability measures for conventional deep learning models. About 10 models whose architecture is ‘explicitly defined’ by domain knowledge are included. However, the relative merits of these architectures or their underlying assumptions are not critically evaluated. Dhillon *et al*. [53] offered a more general systematic review of ML methods for multi-omics-based biomarker identification— again focussing on cancer—without a specific focus on biologically-informed models like VNNs. Abbasi *et al*. [56] reviewed a large number of methods for survival analysis in multi-omics, using deep learning, traditional ML and statistical methods, but not any methods for knowledge integration. A recent systematic review paper by van Hilten *et al*. [59] includes 24 VNN papers, representing probably the most complete survey to date. They compare the popularity of VNNs with other methods, noting the sparsity of VNNs allows handling a greater number of input features than computationally intensive transformer models, though conventional NNs remain the most popular approach. However, the review does not distinguish different VNN architectures or discuss specific design decisions.

### Benchmarks

Leng *et al*. [51] performed a benchmark of deep learning models for multi-omics data fusion in cancer, comparing 6 NNs (including two CNNs and two GNNs), four autoencoders and six variational autoencoders on simulated and real cancer datasets for supervised and unsupervised learning tasks. However, their analysis did not cover any NNs with biologically-informed architectures. Similarly, Hauptmann & Kramer [54] evaluated various deep learning-based multi-modal fusion approaches in drug response prediction and proposed a new method called Omics Stacking, but did not explore any VNNs. Other works extending specific methods include benchmarks of one or more VNN approaches [e.g. 63]: we discuss these in section 5.

### The gap we fill

In summary, most existing reviews either focus broadly on multi-omics ML approaches or are limited to specific contexts such as cancer or general interpretability methods. Our systematic review addresses this gap by focussing specifically on *visible neural network* architectures. We aim to provide a more detailed analysis of technical implementations, discussing the topic of robustness and reproducibility, covering the latest works in this area. Up-to-date coverage includes recent papers from 2024, focussing on applications beyond oncology and with an examination of ways in which VNNs have been both designed, implemented and evaluated, summarizing comparisons with traditional models and with each other.

## 4 Methods

Our aim was to compare architectural design considerations in Binn-like models and identify key contributions in the field. Thus we sought papers—for which the full text was available, to understand the model structure—using NN models informed by biological ontologies, applied to tabular (multi-)omics datasets.

The systematic review was conducted following the Preferred Reporting Items for Systematic Reviews and Meta-Analyses [Prisma; 64] guidelines to ensure transparency and rigour. The Prisma flow diagram (Figure 2) outlines the process of study identification, screening, eligibility assessment and inclusion.

**Figure 2.**
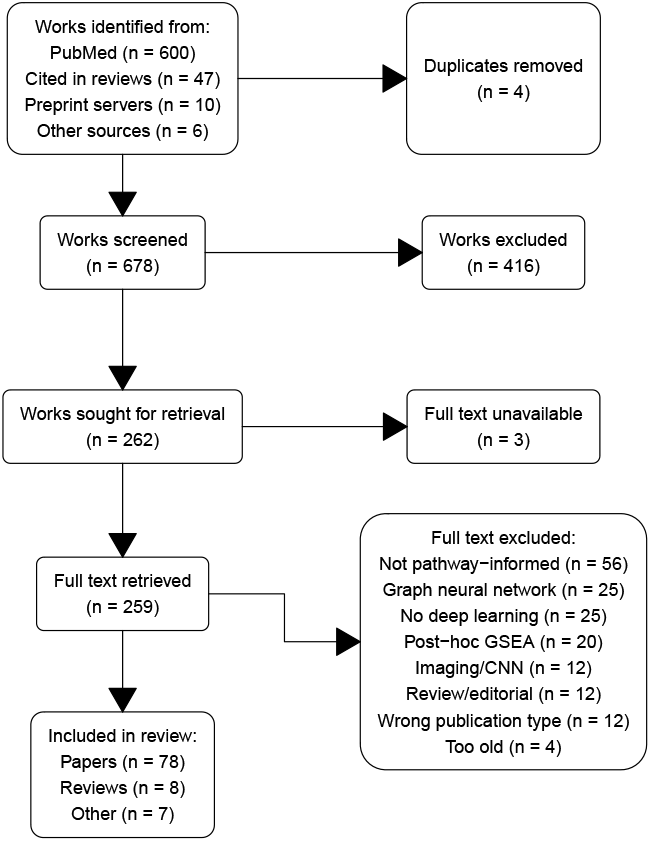
Prisma diagram summarizing article selection

### Search strategy

We conducted a comprehensive search using PubMed to identify relevant papers. Unlike Wysocka *et al*. [43] we chose not to restrict our search to works in oncology, instead focussing on biologically informed models in any application with the query multi-omics AND (deep learning OR computer science OR neural networks OR network analysis OR machine learning) AND (biologically informed), filtered to works published in the period 2018–2024.

However, non-standard terminology in this subfield means this query alone may not capture all relevant papers. For example, Ma *et al*. [65] was one of the first papers to use the term ‘visible neural network’, but ‘multi-omics’ or ‘omic’ does not appear anywhere in the article, nor do phrases like ‘biologically informed’ or ‘pathway’. Other relevant works may be published in conferences or periodicals not indexed by PubMed. Rather than contorting our search query to detect these (admittedly significant) edge cases, we opted to augment our main query with a Google Scholar search for ‘visible neural networks’ as well as adding selected papers manually, including 47 works already included in existing reviews.

Periodically, our database of results has been updated by adding new articles reported as citing key review papers, according to Google Scholar. In this way we were able to include several very recently published works, e.g. Meirer *et al*. [66] and Liu *et al*. [67].

### Screening

After removing duplicates, a total of 678 unique records were screened by title and abstract using inclusion and exclusion criteria as defined above. The screening was performed by three reviewers—with backgrounds in statistics, computer science and bioinformatics— working independently. Disagreements were resolved by majority vote.

### Eligibility

Full-text versions of 262 articles were assessed for eligibility against the predefined criteria, which included methodological journal or conference papers proposing neural network approaches to model omics data that integrate prior biological knowledge into their network architecture. Studies were excluded at this stage if they did not involve NNs, if they were fully connected, non-biologically informed structures, or if biological interpretations only took the form of *post-hoc* gene-set enrichment analysis (GSEA). We focussed our attention on sparsely-connected feedforward neural networks (FFNNs) and autoencoders, while convolutional, attention or graph-based models were generally excluded, as were those designed solely for genomic/proteomic sequences or imaging data.

### Inclusion

86 papers met the eligibility criteria and were included in the final analysis. These studies represent a variety of papers proposing or evaluating biologically-informed neural network architectures, as well as several relevant survey papers (see section 3). We also highlight 7 ‘honourable mentions’: papers that do not strictly meet the inclusion criteria but are interesting examples of alternative approaches.

### Reporting

The full list of included papers is given in Table A.1. The Prisma checklist is provided in the supplementary materials.

### Bibliometrics

Citation counts and relationships were retrieved from the CrossRef API using the R package **rcrossref** [68] to construct a citation network between the papers (see Figure 3). We verified the existence of individual works in PubMed using the NCBI eUtils API [69].

**Figure 3.**
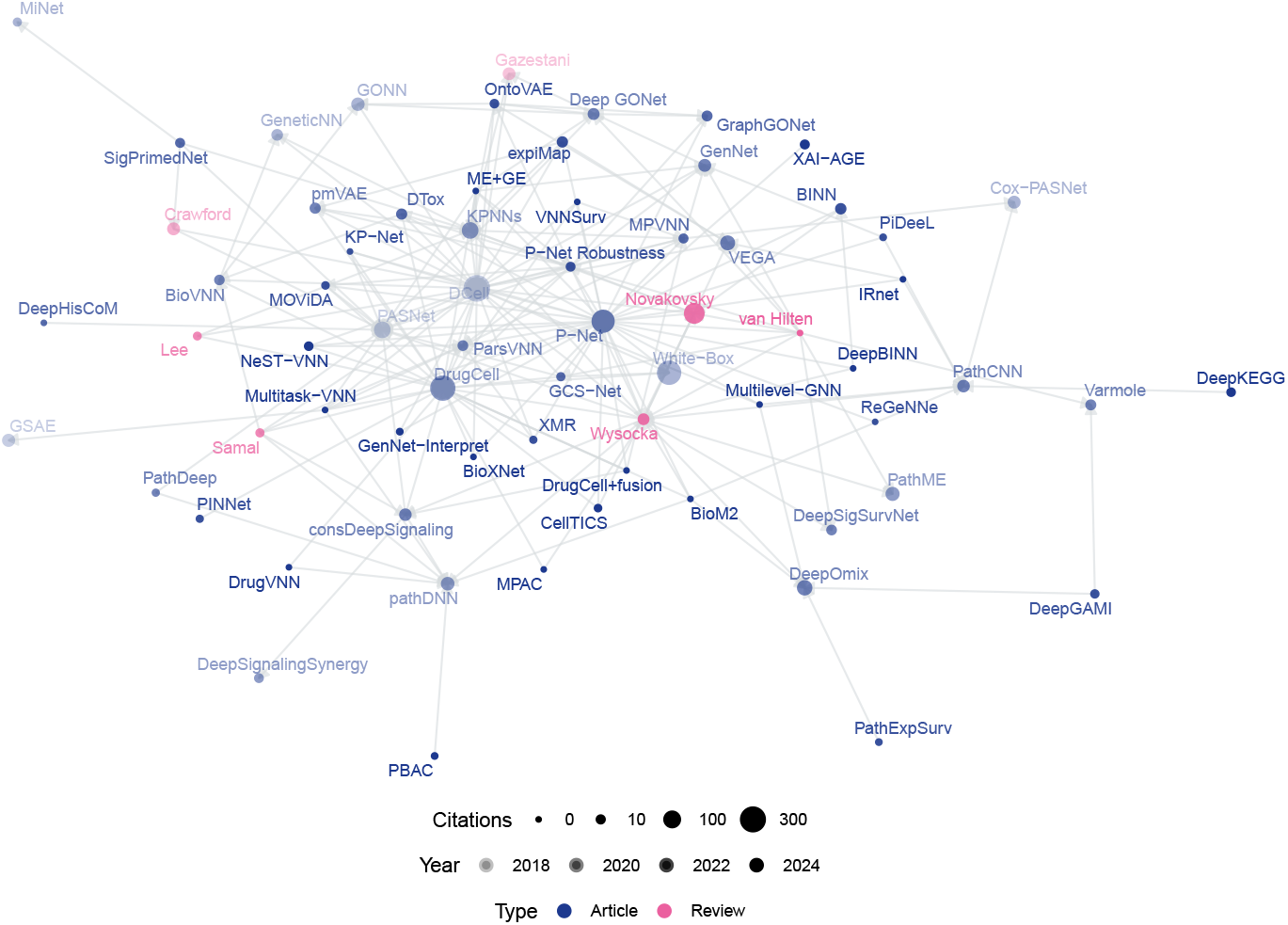
Bibliometric network of VNN papers, scaled by number of citations and shaded by year of publication

## 5 Results

### 5.1 Overview

Our review examines the key advancements in visual neural networks (VNNs), focussing on taxonomy, architectural innovations, methods for evaluation and practical applications. The review identified major developments across diverse architectures—feed-forward neural networks, autoencoders and graph neural networks—evaluated on a variety of omics datasets and knowledgebases. A critical theme is the tradeoff between sparsity for interpretability and the robustness of explanations in complex biological datasets. We also explore how specific design decisions, such as data integration strategies and pathway representations, influence model performance.

We consider ‘strong’ biological guidance [35] where models are constrained by prior knowledge, specifically the ‘in-processing’ paradigm [43] where such information informs the model’s internal architecture. By contrast, wrangling data into a graph structure and applying non-dedicated ML algorithms (such as GNNs) constitutes biologically-informed ‘data pre-processing’ [43] and is not our focus here; similarly we do not dwell on *post-hoc* model explanations or enrichment analyses for non-biologically-informed models [e.g. 33].

Post-2020, VNN research has grown significantly, with key works like DCell [65], DrugCell [70], and P-Net [41] central to the citation network (Figure 3). P-Net has been reproduced by Pedersen *et al*. [71] and built upon by Hartman *et al*. [40], Prosz *et al*. [72], Hartman [73], Hao *et al*. [74] and Hu *et al*. [75], applying the framework to other modalities and targets. Similarly, MOViDA [76] builds upon DrugCell, itself an extension of DCell. Approximately half of recent studies are uncited by prior reviews (see Figure 2), indicating untapped contributions. A full list of retrieved works and associated abbreviations is given in Table A.1.

### 5.2 Taxonomy of architectures

Table 2 divides works according to their broad architecture: feed-forward neural networks (Figure 1d) and autoencoders (Figure 1b) constitute the majority of VNNs, but there also exist some GNNs and CNNs with apparent pathway-aware properties, and other notable approaches.

**Table 2.** Model architectures used in included papers.

| Architecture | *n* | Papers |
| --- | --- | --- |
| FFNN | 53 | BiGMLVQ, BINN, BioM2, BioVNN, BioXNet, Cancer-Net, CellTICS, consDeepSignaling, Cox-PASNet, DCell, Deep GONet, DeepBINN, DeepGAMI, DeepHisCoM, DeepKEGG, DeepOmix, DeepSignalingSynergy, DeepSigSurvNet, DrugCell, DrugCell+ fusion, DrugVNN, DTox, GCS-Net, GeneticNN, GenNet, GenNet-Interpret, GONN, k-DNN, KP-Net, KPNNs, ME+GE, MiNet, MOVIDA, MPVNN, Multitask-VNN, NeST-VNN, P-Net, P-Net Robustness, PAGE-Net, ParsVNN, PASNet, PathDeep, pathDNN, PathExpSurv, PiDeeL, PINNet, Post-hoc node enrichment, SigPrimedNet, Varmole, VNNSurv, White-Box, XAI-AGE, XMR |
| Autoencoder | 10 | ACSNI, expiMap, GONN, GSAE, OntoVAE, PAAE, PathME, pmVAE, priorVAE, VEGA |
| CNN | 5 | DeepGO, PathCNN, Pathway-centric LT, PBAC, ReGeNNe |
| GNN | 5 | DrugVNN, GraphGONet, GSNNs, IRnet, Multilevel-GNN |
| Factor Graph | 2 | FGNN, MPAC |
| Other | 2 | AKLIMATE, PLATYPUS |
| GAN | 1 | omicsGAN |

A VNN is a neural network model with at least one layer of structural sparsity based on connections not directly observable in the main data. (Hence, patient similarity networks [77, Figure 1c] are not VNNs, because their inter-node connections are derived from the data.)

Figure 4 gives a prototypical illustration, with various features of VNN models, described in detail later in this section and tabulated in Table 3. From left to right (input(s) to output) the model’s abstraction becomes progressively more complex as later levels in the neural network correspond to higher tiers in the pathway hierarchy. Such a model can be constructed from a FFNN, with numbers of hidden layers and nodes within each layer corresponding to the desired gene and pathway levels. Initially densely connected, a masking matrix removes inter-layer connections that do not correspond to known biological relations.

**Table 3.** Architectural features in included papers.

| Feature | Value | *n* | Examples |
| --- | --- | --- | --- |
| Pathway Depth | Deep | 29 | BINN, BioXNet, Cancer-Net, CellTICS, Deep GONet, DrugCell, DrugCell+fusion, DTox, FGNN, GCS-Net, GeneticNN, GenNet, GenNet-Interpret, GSNNs, IRnet, k-DNN, KP-Net, KPNNs, ME+GE, MOVIDA, Multitask-VNN, NeST-VNN, OntoVAE, P-Net, P-Net Robustness, Pars-VNN, VNNSurv, XAI-AGE, XMR |
|  | Shallow | 40 | ACSNI, BiGMLVQ, BioM2, BioVNN, consDeepSignaling, Cox-PASNet, DCell, DeepBINN, DeepGAMI, DeepHisCoM, DeepKEGG, DeepOmix, DeepSignalingSynergy, DeepSigSurvNet, DrugVNN, expiMap, GONN, GraphGONet, GSAE, MiNet, MPVNN, Multilevel-GNN, PAAE, PAGE-Net, PASNet, PathCNN, PathDeep, pathDNN, PathExpSurv, PathME, PBAC, PiDeeL, PINNet, pmVAE, priorVAE, ReGeNNe, SigPrimedNet, Varmole, VEGA, White-Box |
|  | NA | 1 | MPAC |
| Multi-omics | Multi-omics | 29 | BiGMLVQ, BioM2, BioVNN, BioXNet, Cancer-Net, consDeepSignaling, DeepGAMI, DeepHisCoM, DeepKEGG, DeepOmix, DeepSignalingSynergy, DeepSigSurvNet, DrugVNN, GCS-Net, GSNNs, KP-Net, ME+GE, MiNet, MOVIDA, MPAC, Multilevel-GNN, Multitask-VNN, NeST-VNN, P-Net, P-Net Robustness, PathCNN, PathME, Varmole, VNNSurv |
| Hidden Layers | Yes | 35 | BiGMLVQ, consDeepSignaling, Cox-PASNet, DeepBINN, DeepGAMI, DeepHisCoM, DeepOmix, DeepSignalingSynergy, DeepSigSurvNet, DrugCell+fusion, expiMap, GCS-Net, GONN, GSAE, GSNNs, IRnet, MiNet, MOVIDA, Multilevel-GNN, OntoVAE, PAAE, PAGE-Net, PAS-Net, PathCNN, PathDeep, pathDNN, PBAC, PiDeeL, PINNet, pmVAE, priorVAE, SigPrimedNet, Varmole, VEGA, VNNSurv |
| Mixed Layers | Yes | 4 | BioM2, expiMap, PAGE-Net, PINNet |

**Figure 4.**
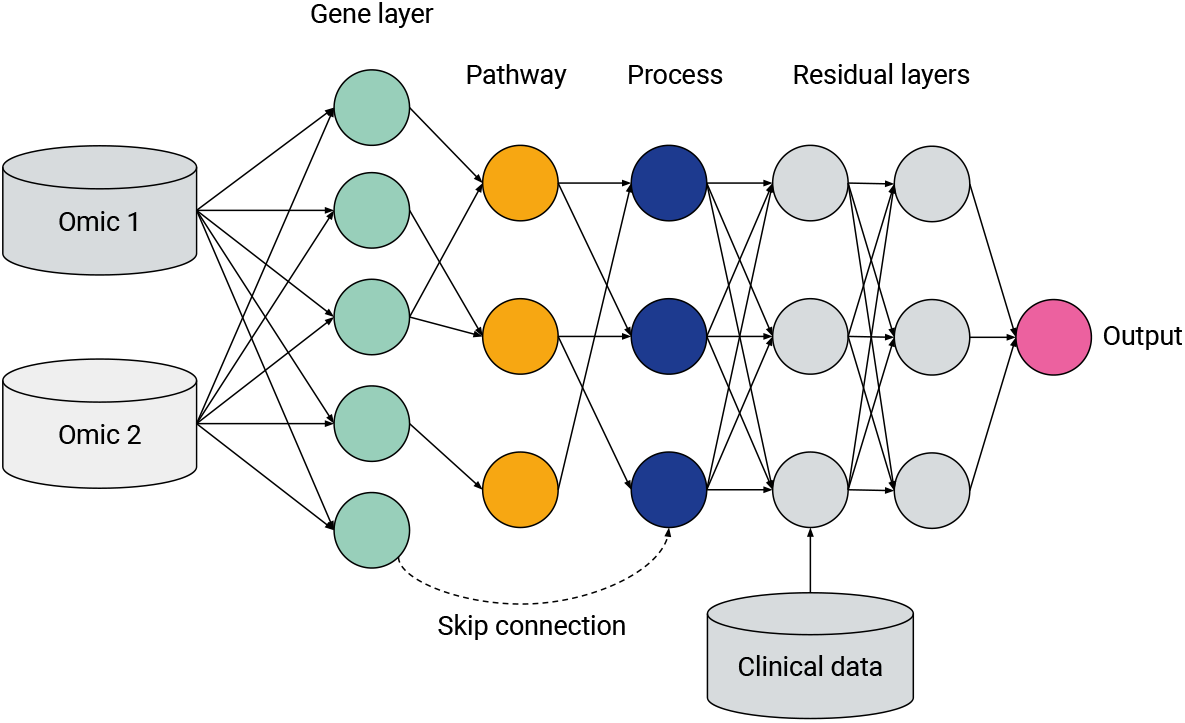
General hierarchical structure of a visible neural network, featuring multi-omics integration, a hierarchy of genes, multilevel pathways or biological processes (green, yellow and blue, respectively), a ragged structure with skip connections and layers containing fully connected ‘residual’ nodes. Data not mappable to pathways can be included via intermediate or late fusion

#### Feed-forward networks

A ‘standard’ NN for supervised learning, FFNNs may be applied to classification, regression or survival analysis tasks. Feed-forward VNNs may have deep structures with multiple layers of nested pathways, such as DCell [65] and P-Net [41]; however the simplest networks, e.g. Cox-PASNet [61] include just one pathway layer.

#### Autoencoders

Autoencoders are a class of deep learning models for unsupervised learning and consist of three components: an encoder, a latent space and a decoder (see Figure 1b). Input features are encoded through one or more hidden layers into a ‘bottleneck’ layer representing a low-dimensional latent space; a decoder, typically mirroring the architecture of the encoder, then reconstructs the input features from this space. Some of the earliest Binns are autoencoders, e.g. GSAE [78]. A typically used extension of this idea are variational autoencoders (VAE) which are probabilistic and generative models learning a multivariate distribution (e.g. Gaussian) as latent space, enabling latent space disentanglement and compactness. Examples include Vega [79] and ExpiMap [80]; other models come in both AE and VAE variants, e.g. PAAE/PAVAE [81], or FFNN and AE variants, e.g. GONN/GOAE [82]. Liu *et al*. [83] proposed ‘pathway-informed priors’ where knowledge is integrated into the VAE loss term.

#### Convolutional neural networks

CNNs have been widely adopted in the analysis of structured datasets like images and sequences. Applying CNNs to multi-omics data is possible by combining with other data modalities such as sequences [DeepGo; 84] or by synthesizing ‘pathway images’ from tabular data [PathCNN; 85].

#### Graph neural networks

GNNs can directly leverage the topological structure of biological data, making them particularly well-suited for tasks where the relationships between entities need to be explored. For example, GraphGONet [86] uses the structure of the Gene Ontology graph directly; Yan *et al*. [87] used a GNN to embed genes, followed by a ‘pathway aggregation block’ resembling a FFNN, to obtain pathway-level features. As GNNs operate primarily through learned graph representations, they do not inherently enforce predefined biological hierarchies, which can make them less suitable for applications where pathway-level interpretability and strict adherence to known biological relationships are necessary.

Some authors adopt hybrid approaches, for example: ReGeNNe [88] combines GNN and CNN layers; GraphGONet [86] claims to offer advantages of an FFNN and a GNN; DeepKegg [89] introduces a ‘pathway self-attention module’. Biologically informed generalized matrix learning vector quantization [BiGMLVQ; 42] bills itself as a non-deep-learning approach but is nevertheless NN based. BioM2 [90] performs pathway-level feature selection, concatenating features with filtered inputs unmapped to pathways. While this approach can be performed in a deep learning pipeline, the authors note it could also be performed using multi-stage logistic regression or other machine learning methods. In a more network focused approach, Kim *et al*. [91] and Kim *et al*. [92] employ a random walk algorithm across an integrated pathway network based on the Kyoto Encyclopædia of Genes and Genomes (Kegg) pathways to derive pathway-level scores for downstream tasks such as survival prediction. However, this approach does not directly incorporate a biologically informed neural network structure. Uzunangelov *et al*. [93] present Aklimate, an example of a stacked kernel learner incorporating pathway knowledge.

### 5.3 Data and applications

Figure 5 shows gene expression (transcriptomics/mRNA) is the dominant omics modality, featuring in more than half of papers, followed by copy number variations (CNV); DNA mutations, including single nucleotide polymorphisms (SNPs), were also a common data type. As shown in Table 3, 29 models integrated multi-omics data, with some models such as P-Net [41] ostensibly applicable to any number and combination of omics levels (in the paper itself they used just two). A limited number of models—especially for survival analysis—integrated omics or multi-omics data with ‘non-omics’ inputs not mappable to pathways: Cox-PASNet [61] and MiNet [94], combine gene expression, DNA methylation and copy number variations with clinical data, such as patient age, via late fusion. DrugCell [70, 95] extended DCell [65] by integrating mutation data with embeddings of the chemical structure of different drugs, also via late fusion; however later analysis by Nguyen *et al*. [96] found mid-or early fusion models performed better. Other examples of such drug–genotype modular networks include BioXNet [97], MOViDA [76] and XMR [98].

**Figure 5.**
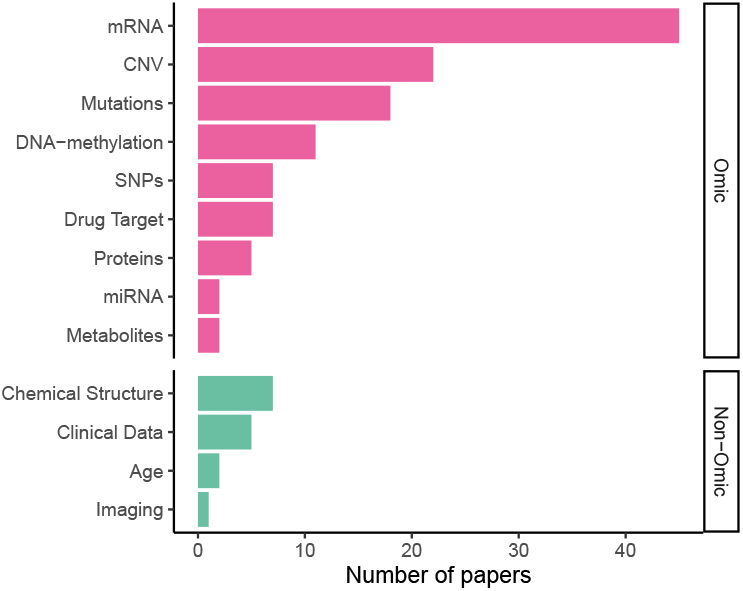
Input data types in published VNN models

As shown in Table A.2, oncology is a major application area for VNNs, with 46 papers; 13 were concerned with drug response [e.g. consDeepSignaling; 99]; 4 with cellular processes; 3 with schizophrenia and 3 with Covid-19. While this shows cancer research is significant, VNNs are not limited to this disease.

Many authors used data from common sources, with omics data from the Cancer Genome Atlas (TCGA) and Gene Expression Omnibus (GEO) particularly popular; see Figure 6. Some authors used their own collected data or else did not clearly specify an existing database as their source. Use of common data sources may make replication of results more straightforward (nearly every paper provided analysis code: Table A.4) but may also raise questions about their real-world generalizability, given sampling biases present in these cohorts. (Kegg) and Reactome were equally commonly chosen as sources of biological ontology information, followed by Gene Ontology and MSigDB. All are publicly accessible databases. Kim & Lee [100] highlighted differences in performance for the downstream task depending on the ontology used.

**Figure 6.**
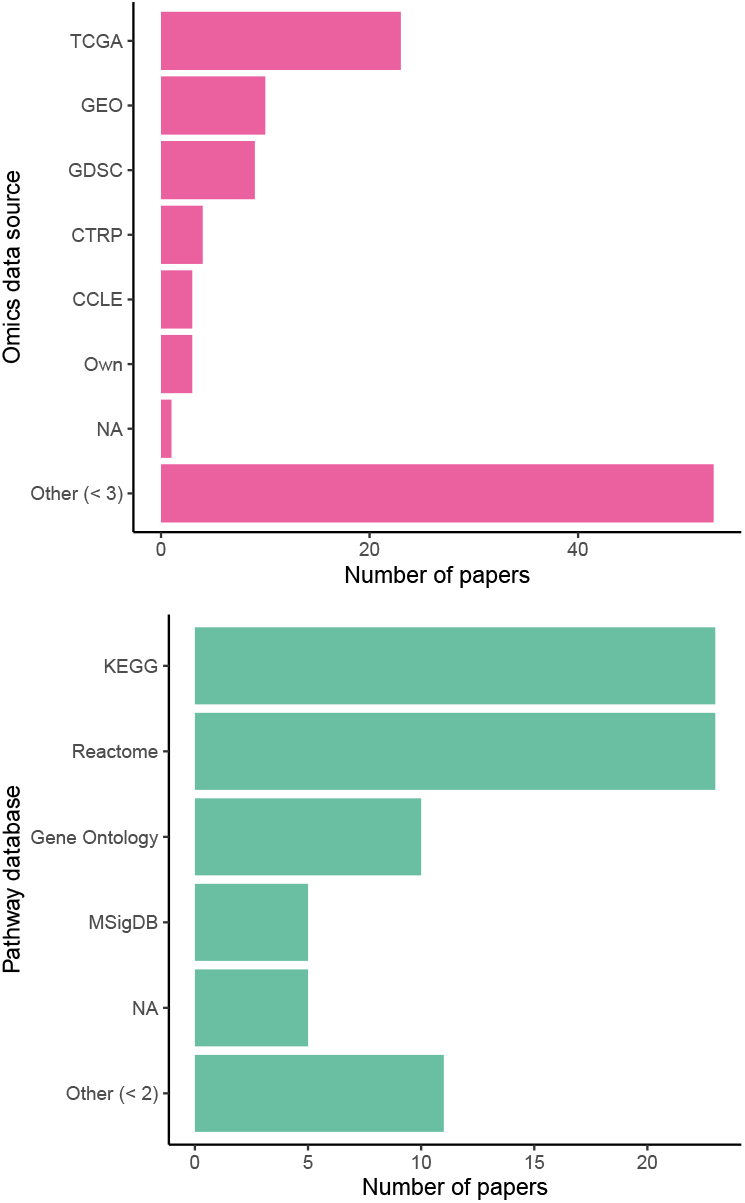
Sources of omics data and pathway knowledge in biologically-informed models

### 5.4 Evaluating biologically informed architectures

Assessing the usability of VNNs requires comprehensive evaluation of predictive performance, interpretations and robustness against appropriate baselines. Of included papers, 56 compared with traditional (non-NN) models and 54 compared with non-biologically-informed NNs, with just 22 comparing their proposed models with other Binn variants, meaning most VNNs are evaluated in isolation to other biologically-informed models.

#### 5.4.1 VNNs perform similarly to denser models

Fortelny & Bock [34] posited that Binns operate as ‘information processing units’ using network-based computations to regulate their states. Thus, by mirroring cellular regulatory systems, VNNs provide a deeper understanding of these mechanisms, and can offer similar prediction performance to ‘black box’ fully-connected networks while being significantly more sparse. However, redundancy in the structure of VNNs leads to variability in edge weights, making explanations less robust—a problem that may be mitigated via dropout on hidden nodes during training. Similarly, Huang *et al*. [101] argued that redundancy among nodes and edges can lead to overfitting and less robust explanations; they proposed a pruning mechanism to mitigate the issue, resulting in model simpler than DrugCell while offering superior accuracy.

Kuenzi *et al*. [70] compared their VNN to a ‘black-box’ neural network with equivalent depth and sparsity, finding broadly similar performance and superior accuracy to a non-deep-learning model. In contrast, Pedersen *et al*. [71] compared P-Net with a randomly-connected network. They observed a clear decrease in performance for the random connections in comparison to the Binn. Lin *et al*. [102] also found biologically-sparse networks outperformed fully-connected models. However a fully-connected network with random masked nodes is not necessarily optimal for multi-omics if it assumes early fusion [54]; VNNs have been shown sensitive to fusion stage [96], so but early-integrated baselines could be ‘strawmen’.

#### 5.4.2 Performance is sensitive to architecture

Several studies have compared different biologically informed architectures on the same data to assess their relative benefits. For instance, PINNet [100] was compared against traditional ML models and dense NNs. They also compared two types of VNNs, informed by GO and Kegg respectively, which outperformed the non-biologically-informed models. Meanwhile, PBAC [103] conducted an ablation study on its architecture, revealing that removing the biological information mask and the attention layer, respectively, reduced the performance of drug response prediction. However, they did not benchmark their architecture against other models.

#### 5.4.3 Robustness of interpretations

Since VNNs rely on the same biological knowledge as commonly used gene-or pathway-enrichment analysis frameworks, they inherit similar problems, such as balancing between specific pathways with few genes and broad pathways with many genes. Additionally, VNNs can be prone to overfitting and instability resulting from small sample sizes, leading to significant changes with different train-test splits or initializations [63]. Common strategies for the training of VNNs are the introduction of dropout layers to stabilize training, in particular when overlapping ontology terms are used. Further, weights can be restricted e.g. by weight decay configuration or by restricting the direction (positive weight) to fix direction. Additionally, Esser-Skala & Fortelny [63] assessed the robustness of interpretations in several models, including DTox and P-Net [36, 41].

Meirer *et al*. [66] raised the question of robustness of biomarker signatures found by Binns. Their solution, DeepBinn, fits a sub-network per pathway—each a NN with a fixed number of hidden layers—whose output weight measures that pathway’s importance. By comparing the ranks of pathways over successive initializations, the authors claim to yield a ‘robust’ signature.

### 5.5 Architectural design

Even within the taxonomies listed in Table 2, VNN architectures differ considerably, based on different design considerations highlighted in Figure 4.

#### 5.5.1 Pathway nodes

Some VNN architectures have multiple hidden nodes per gene or pathway, representing their ability to perform multiple tasks concurrently. For example, DCell [65] and derivatives [70, DrugCell, 76, MOViDA] have 6 nodes per pathway; OntoVAE [104] has 3. However, most other architectures map each entity to at most one node in the network, simplifying the representation at the possible expense of predictive power. Perhaps the most crucial design choice is the selection of the source and type of biological knowledge to be incorporated in a VNN. However, surprisingly little efforts have been made in the field to analyse and study the impact on choosing Reactome pathways, KEGG, GO-terms or other sources. Since these sources all have their strength and weaknesses in relation to coverage (number of genes represented), overlap between terms or curation quality, a big impact on VNNs and their architectural choices is expected. For example for VAEs authors state that overlap of genes across pathways is countered by dropout layers [79], similarly reported in feed/forward VNNs by Fortelny & Bock [34]. ParsVNN [101] approaches the problem of redundant subsystems in VNN models by a pruning mechanism.

Lastly, the chosen database or ontology, as well as the level of hierarchy, ideally matches the purpose of the model e.g. using KEGG metabolic networks for insights into cancer cell metabolism or Reactome immune pathways for studying infections. Currently, most studies using VNNs are explorative and typically do not restrict incorporated biological knowledge in a purpose-driven way. However, PathExpSurv [62] uses a ‘pathway expansion’ phase to adaptively adjust pathway-gene connections, potentially enhancing interpretability. Also expanding on the idea of including pathway information, MPVNN [105] is uses mutation assays to build multiple (mutated) versions of a given pathway.

#### 5.5.2 Auxiliary layers (layer-wise loss)

DCell also introduced ‘auxiliary layers’, a method for intermediate pathway layers (for models who have them) to contribute to the loss function and to allow attribution to particular hierarchical levels. P-Net adopts a similar approach of adding ‘predictive layers’ with sigmoid activation after each hidden layer, with later layers weighted more highly in the loss function.

#### 5.5.3 Hidden and mixed layers

Some architectures include fully-connected hidden layers that are not *a priori* biologically interpretable. These layers introduce additional complexity to the network, potentially improving predictive power but at the expense of interpretability. For example, models like PASNet [74] and MiNet [94] include this structure. However, the rationale behind including such layers is not always clearly explained.

The inclusion of these hidden layers may allow the model to capture latent interactions not accounted for by existing biological knowledge, but this assumption requires further validation.

Other architectures include fully-connected nodes in the same layer as the interpretable nodes, a design we will call ‘mixed layers’; with the the additional nodes denoted ‘residual’ nodes for their ostensible aid to model performance or simply ‘hidden’ nodes due to their lack of biological interpretability.

In most works the authors do not mention why they chose a given design. However, some works have performed ablation studies with and without hidden layers or nodes in the architecture: Lin *et al*. [102] compare multiple dense NNs and VNNs with one or two fully connected hidden layers, seeing the best overall performance in the networks with two additional hidden layers in a clustering task. They also found pretraining not beneficial in comparison to training from scratch for clustering cells, but it increased performance for cell type retrieval (classification task). In contrast in [42, BiGMLVQ] the model with more prototype layers (classification layers) performed worse, which the authors attributed to overfitting. Fortelny & Bock [34] showed that drop-out on pathway level, as well as input layers, increased robustness to redundancies and imbalances inherent to biological networks.

In particular, autoencoders using biological knowledge to guide representation learning rely heavily on the combination of fully-connected encoders and sparse decoders representing pathway or ontology relationships. However, only authors of VEGA [79] studied other possible architectures and combinations while others like expiMap [80] or OntoVAE [38] adopted the same general structure and a theoretical analysis or broad benchmark of different encoder-decoder combinations has not been done to our knowledge. Neither are we aware of any work that performed a systematic architecture search with the pathway layers or additional fully connected layers on feed-forward VNNs.

#### 5.5.4 Shallow and deep hierarchies

Biologically-informed sparse neural network architectures involve a kind of inductive bias, wherein the hidden layers represent biological entities or processes and the connections between the layers represent biological relations. In principle, such a network can be constructed by starting with a fully-connected network and then applying a masking matrix corresponding to the existence or strength of relations between the biological entities. However, given heterogeneous pathway databases and data processing pipelines, the method of converting a hierarchy of biological pathways into a neural network is by no means standardized. For example, a pathway may contain genes and other pathways: should the network include skip connections to account for this (see Figure 4)? How many hidden layers or nodes in each layer should be chosen before certain paths are ‘pruned’ from the model?

Many architectures incorporate a single layer for pathways following the gene layer, but others extend this to multiple levels to represent a hierarchy of pathways. The methods for constructing these hierarchical relationships vary across models, and often it is not explicitly clear how a ‘top-level’ pathway or biological process is defined. For instance, decisions on whether intermediate pathways should be merged or truncated are typically not well-documented. Ma & Zhang [106] proposed factor graph neural networks (FGNNs), which include a single pathway layer, but may be ‘unrolled’ to a deeper structure to add greater expressive power.

The early pathway-guided neural network architectures are rather complex, incorporating deep pathway hierarchies. ParsVNN [101] applied proximal alternative linearized minimization to the NP-hard problem of *l*_0_ norm and group lasso regularization in order to prune redundant edges between pathways, resulting in simpler structures.

Other authors argue for the simplicity and interpretability of a shallow network [42, 78] with just one gene layer aggregating inputs from one or more omics levels, followed by a single pathway layer.

One technical consideration when building a deeper network is how to handle pathways that are not all the same depth of hierarchy. The bottom of a pathway ontology is not well defined, so networks such as P-Net and later Cancer-Net [71] start from the most abstract level of ‘biological processes’, defined as Reactome pathways with no parents, successively adding layers for each generation of child pathways, stopping at a user-selected depth, for example, 6 layers. If a branch of the Reactome tree does not have so many levels, then ‘dummy nodes’ are added to allow implementation in a layered machine learning framework like PyTorch. However, since each of these dummy nodes has a nonlinear activation, some information is inevitably lost or altered. In contrast, DrugCell [70] employs direct skip connections, which may be more challenging to implement in common ML frameworks but offer greater flexibility in modelling complex biological relationships.

The notion of skip connections can be extended further. Whereas a VNN is typically formulated as a directed acyclic graph with discrete sequential layers, Evans *et al*. [107] recently proposed ‘graph-structured neural networks’ (GSNNs)—not to be confused with GNNs— an extension of VNNs that allows for cycles and self-loops. Like GNNs, GSNNs use an input graph as an inductive bias to constrain the information flow in the neural network. Unlike GNNs, GSNNs do not share weights across nodes; instead, each node is associated with its own distinct neural network.

### 5.6 Scientific discovery

VNNs being more performant than their densely-connected counterparts raises the possibility of discovering pathway structures through regularization. That is, if a VNN really is better than an equivalently sparse model, then neural architecture search, for example via pruning, could in principle learn such a structure without prior knowledge integration [108]. Such an approach to scientific knowledge discovery—which would enable prediction of new pathway relations, akin to knowledge graph edge prediction—has not yet been widely explored in the VNN literature.

Though ParsVNN [101] prunes irrelevant pathways, it starts with a biologically-informed structure. Similarly DeepHisCoM [109] and DeepBinn [66] explore nonlinear relationships only among existing pathways. PathExpSurv [62] extended pathway knowledge integration by performing ‘pathway expansion’ to include pathways that may not exist in the original database. First a biologically-sparse network is trained, then fine-tuned with dense connections; weights that are not regularized in the second stage may be indicative of undiscovered pathways. Mixed layers (see Table 3), containing both interpretable and non-interpretable nodes, might allow models to harness predictive performance beyond the constraints of the pathway database, but so far this has not been applied to discovery of new pathways.

### 5.7 Resources and tools

Most papers introducing methods provide open-source code for reproducibility (Table A.4), however, the reusability and maintainability of some of these code-bases is not necessarily guaranteed, especially if raw data or data preprocessing code are not provided or the repository hard-coded for a specific dataset or file system. Nonetheless, Pedersen *et al*. [71] was able to reproduce the results of P-Net [41], updating the code to newer ML frameworks. van Hilten *et al*. [110] released their GenNet framework, which has since been extended with modules for interpretability [111].

A recently developed package called binn [40], focussed on proteomics, offers the capability to build VNNs with a given input pathway set. Autoencodix [39] is a framework for building different autoencoder architectures, which includes one ontology-based architecture as an option.

There remains a gap, however, for a user-friendly, general-purpose package that supports multi-omics data inputs and different pathway databases and allows exploration of different design decisions to ensure the robustness of the yielded predictions and explanations. Especially interesting would be a highly general package capable of modelling non-biological ontologies (such as in chemistry or social sciences) using a sparse VNN framework.

## 6 Discussion

VNNs have seen increasing adoption since their emergence around 2017–18. Their versatility is evidence in applications spanning protein classification, survival analysis, diagnosis and drug-interaction prediction [61, 65, 74, 84]. Compared to full-connected neural networks and other ML algorithms, VNNs often demonstrate comparable or superior performance. Studies such as Pedersen *et al*. [71] highlight that randomized sparse networks of comparable size underperform in comparison, underscoring the value of integrating biological knowledge. This integration enhances neural networks’ ability to extract signals from relatively small and tabular datasets. However, many studies fail to benchmark against traditional ML methods, which can outperform neural networks [112], do not compare with non-biologically-informed NNs to quantify the value of knowledge integration, or do not compare their implementation with existing VNN frameworks. Interpretability is a key advantage of VNNs over ‘black box’ dense neural networks. By constraining networks with biological pathways, VNNs reduce the function space, trading universality for an inductive bias that promotes faster convergence and enables pathway-level insights via node activations [113]. Despite this potential, further theoretical exploration of these inductive biases is required. Emerging evidence suggests that node activations in VNNs can be mapped to their biological counterparts to yield novel insights verifiable in the laboratory [41], but no studies have conclusively shown that these activations are both data-driven and shaped by the inductive bias. Sparse network randomizations studies remain insufficient to fully elucidate the function spaces of these models. Additionall,y the reliance on simplified gene-to-pathway mappings often neglects directional and regulatory relationships, limiting real-world applicability. Flexible architectures with less restrictive activation functions may better capture the multi-layered complexity of cellular biology, bridging trends in explainability and the training of molecular foundation models [114–116].

Another limitation lies in the completeness and quality of pathway databases. While these resources are invaluable, they provide an incomplete picture of cellular processes, particularly for non-human or non-model organisms where database curation is often automated and less robust. Only a few studies, such as Hou *et al*. [62] and Park *et al*. [109], have explored the use of VNNs for pathway expansion or the discovery of new biological relationships, leaving this area largely untapped.

Similarly, the impact of different biological databases or knowledge graphs on performance remains underexplored. Variations in database focus and quality could significantly influence VNN results, but most studies do not benchmark their models across multiple databases. Finally, nearly all VNN architectures rely on early fusion for multi-omics data, which can result in the loss of structural information and diminished performance. For example, Pedersen *et al*. [71] found that fusion strategies had a greater impact on performance than randomised sparse connections. Intermediate fusion techniques, as proposed by Hauptmann & Kramer [54], may offer a more effective approach. However, there remains a gap in developing VNN architectures that incorporate these advanced strategies.

## 7 Conclusion

Visible neural networks (VNNs) represent a promising advancement in multi-omics data integration, providing a unique combination of predictive performance and biological interpretability. This systematic review has highlighted critical trends, including the importance of sparse architectures informed by biological priors, the impact of design choices on model robustness, and the variability in performance across datasets and ontologies. While the field has grown substantially since 2020, challenges remain in standardising terminology, benchmarking methodologies and improving reproducibility.

Future research should prioritise systematic evaluations of VNN architectures, particularly regarding the robustness of pathway-level interpretations and the interplay between sparsity and prediction accuracy. Expanding the use of pathway databases, integrating multi-omics modalities with more flexible fusion strategies and exploring novel applications beyond oncology could further enhance their utility. Additionally, developing general-purpose, user-friendly software frameworks for constructing and evaluating VNNs would foster broader adoption and reproducibility. Discovering novel pathway relations through neural architecture search remains an untapped opportunity. By addressing these challenges, VNNs have the potential to unlock novel biological insights, bridge gaps in multi-omics research and contribute to advances in personalised medicine.

## Acknowledgements

We are grateful for the input of Derian Boer, whose helpful suggestions improved the manuscript. This work was funded by the German Federal Ministry of Education and Research (BMBF), grant number 03ZU1202JA in the frame of Clusters4Future project curATime.

## A Included works

**Table A.1:** Included VNN papers, identifiers and abbreviations.

| Abbreviation | Reference | DOI |
| --- | --- | --- |
| ACSNI | Anene *et al.* [117] | 10.1016/j.patter.2021.100270 |
| BiGMLVQ | Voigt *et al.* [42] | 10.5220/0012420700003657 |
| BINN | Hartman *et al.* [40] | 10.1038/s41467-023-41146-4 |
| BioM2 | Zhang *et al.* [90] | 10.1093/bib/bbae384 |
| BioVNN | Lin & Lichtarge [118] | 10.1093/bioinformatics/btab137 |
| BioXNet | Yang *et al.* [97] | 10.1101/2024.01.29.576766 |
| Cancer-Net | Pedersen *et al.* [71] | 10.48550/arXiv.2309.16645 |
| CellTICS | Yin & Chen [119] | 10.1093/bib/bbad449 |
| consDeepSignaling | Zhang *et al.* [99] | 10.3389/fbinf.2021.639349 |
| Cox-PASNet | Hao *et al.* [120] | 10.1186/s12920-019-0624-2 |
| DCell | Ma *et al.* [65] | 10.1038/nmeth.4627 |
| Deep GONet | Bourgeais *et al.* [121] | 10.1186/s12859-021-04370-7 |
| DeepBINN | Meirer *et al.* [66] | 10.1109/SDS60720.2024.00044 |
| DeepGAMI | Chandrashekar *et al.* [122] | 10.1186/s13073-023-01248-6 |
| DeepHisCoM | Park *et al.* [109] | 10.1093/bib/bbac171 |
| DeepKEGG | Lan *et al.* [89] | 10.1093/bib/bbae185 |
| DeepOmix | Zhao *et al.* [123] | 10.1016/j.csbj.2021.04.067 |
| DeepSignalingSynergy | Zhang *et al.* [99] | 10.1101/2020.04.10.036491 |
| DeepSigSurvNet | Feng *et al.* [124] | 10.1186/s12859-020-03850-6 |
| DrugCell | Kuenzi *et al.* [70] | 10.1016/j.ccell.2020.09.014 |
| DrugCell editorial | Greene & Costello [95] | 10.1016/j.ccell.2020.10.014 |
| DrugCell + fusion | Nguyen *et al.* [96] | 10.1093/bib/bbae227 |
| DrugVNN | Xie *et al.* [125] | 10.1101/2024.02.07.579280 |
| DTox | Hao *et al.* [36] | 10.1016/j.patter.2022.100565 |
| expiMap | Lotfollahi *et al.* [80] | 10.1038/s41556-022-01072-x |
| FGNN | Ma & Zhang [106] | 10.48550/arXiv.1906.00537 |
| GCS-Net | Hu *et al.* [75] | 10.1155/2022/2965166 |
| GeneticNN | Eetemadi & Tagkopoulos [126] | 10.1093/bioinformatics/bty945 |
| GenNet | van Hilten *et al.* [110] | 10.1038/s42003-021-02622-z |
| GenNet-Interpret | van Hilten *et al.* [111] | 10.1101/2024.02.27.582086 |
| GONN | Peng *et al.* [82] | 10.1186/s12859-019-2769-6 |
| GraphGONet | Bourgeais *et al.* [86] | 10.1093/bioinformatics/btac147 |
| GSAE | Chen *et al.* [78] | 10.1186/s12918-018-0642-2 |
| GSNNs | Evans *et al.* [107] | 10.1101/2024.02.28.582164 |
| IRnet | Jiang *et al.* [127] | 10.1016/j.jare.2024.07.036 |
| k-DNN | Ciallella *et al.* [37] | 10.1021/acs.est.1c02656 |
| KP-Net | Zhang *et al.* [128] | 10.1097/CJI.0000000000000475 |
| KPNNs | Fortelny & Bock [34] | 10.1186/s13059-020-02100-5 |
| ME + GE | van Hilten *et al.* [129] | 10.1038/s41540-024-00405-w |
| MiNet | Hao *et al.* [94] | 10.1007/978-3-030-20242-2_10 |
| MOViDA | Ferraro *et al.* [76] | 10.1093/bioinformatics/btad432 |
| MPAC | Liu *et al.* [67] | 10.1101/2024.06.15.599113 |
| MPVNN | Ghosh Roy *et al.* [105] | 10.1093/bioinformatics/btac636 |
| Multilevel-GNN | Yan *et al.* [87] | 10.1093/bib/bbae184 |
| Multitask-VNN | Zhao *et al.* [130] | 10.1158/2159-8290.CD-23-0641 |
| NeST-VNN | Park *et al.* [131] | 10.1038/s43018-024-00740-1 |
| OntoVAE | Doncevic & Herrmann [104] | 10.1093/bioinformatics/btad387 |
| P-Net | Elmarakeby *et al.* [41] | 10.1038/s41586-021-03922-4 |
| P-Net Robustness | Esser-Skala & Fortelny [63] | 10.1038/s41540-023-00310-8 |
| PAAE | Avelar *et al.* [81] | 10.48550/arXiv.2306.05813 |
| PAGE-Net | Hao *et al.* [132] | 10.1142/9789811215636_0032 |
| ParsVNN | Huang *et al.* [101] | 10.1093/nargab/lqab097 |
| PASNet | Hao *et al.* [74] | 10.1186/s12859-018-2500-z |
| PathCNN | Oh *et al.* [85] | 10.1093/bioinformatics/btab285 |
| PathDeep | Park *et al.* [133] | 10.3390/ijms222111531 |
| pathDNN | Deng *et al.* [134] | 10.1021/acs.jcim.0c00331 |
| PathExpSurv | Hou *et al.* [62] | 10.1186/s12859-023-05535-2 |
| PathME | Lemsara *et al.* [135] | 10.1186/s12859-020-3465-2 |
| PBAC | Deng *et al.* [103] | 10.1111/jcmm.18298 |
| PiDeeL | Kaynar *et al.* [136] | 10.1093/bioinformatics/btad684 |
| PINNet | Kim & Lee [100] | 10.3389/fnagi.2023.1126156 |
| pmVAE | Gut *et al.* [137] | 10.1101/2021.01.28.428664 |
| priorVAE | Liu *et al.* [83] | 10.1371/journal.pcbi.1011198 |
| ReGeNNe | Sharma & Xu [88] | 10.1093/bioinformatics/btad679 |
| Review: Crawford | Crawford & Greene [50] | 10.1016/j.copbio.2019.12.021 |
| Review: Gazestani | Gazestani & Lewis [49] | 10.1016/j.coisb.2019.04.001 |
| Review: Lee | Lee & Kim [35] | 10.3345/cep.2021.01438 |
| Review: Novakovsky | Novakovsky *et al.* [55] | 10.1038/s41576-022-00532-2 |
| Review: Samal | Samal *et al.* [52] | 10.3389/fbinf.2022.1036963 |
| Review: van Hilten | van Hilten *et al.* [59] | 10.1093/bib/bbae449 |
| Review: Wysocka | Wysocka *et al.* [43] | 10.1186/s12859-023-05262-8 |
| SigPrimedNet | Gundogdu *et al.* [138] | 10.1186/s13040-021-00285-4 |
| Varmole | Nguyen *et al.* [139] | 10.1093/bioinformatics/btaa866 |
| VEGA | Seninge *et al.* [79] | 10.1038/s41467-021-26017-0 |
| VNNSurv | Tan *et al.* [140] | 10.1016/j.csbj.2024.07.019 |
| White-Box | Yang *et al.* [141] | 10.1016/j.cell.2019.04.016 |
| XAI-AGE | Prosz *et al.* [72] | 10.1038/s41598-023-50495-5 |
| XMR | Wang *et al.* [98] | 10.3389/fbinf.2023.1164482 |

## B Data and applications

**Table A.2:** Disease areas and applications covered in included papers.

| Disease | *n* |
| --- | --- |
| Cancer | 46 |
| Drug Response | 13 |
| Cellular Processes | 4 |
| Schizophrenia | 3 |
| Covid-19 | 3 |
| Other | 15 |
| NA | 8 |

**Table A.3:** Output formats of neural networks in included papers.

| Output format | *n* |
| --- | --- |
| Classification | 38 |
| Regression | 15 |
| Survival | 12 |
| Autoencoder | 10 |
| Review | 7 |
| Survival as Classification | 3 |
| Clustering | 1 |
| NA | 1 |

## C Prisma checklists

The Prisma checklist is available to download (.docx format).

## D Code availability

**Table A.4:**
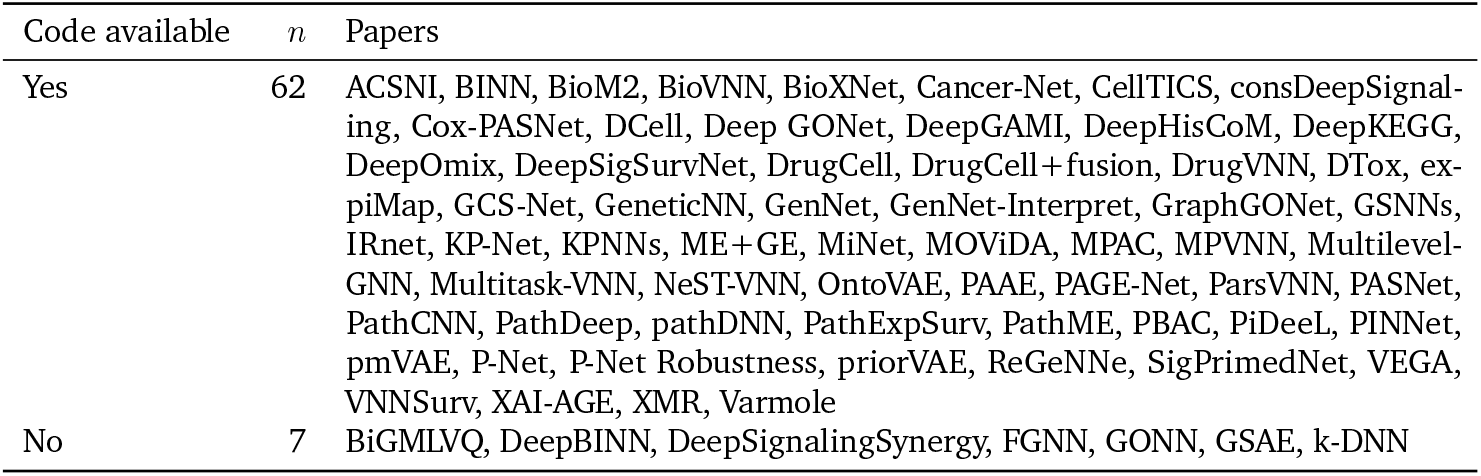
Code availability in papers proposing or applying methods.

| Code available | *n* | Papers |
| --- | --- | --- |
| Yes | 62 | ACSNI, BINN, BioM2, BioVNN, BioXNet, Cancer-Net, CellTICS, consDeepSignal-<br>ing, Cox-PASNet, DCell, Deep GONet, DeepGAMI, DeepHisCoM, DeepKEGG,<br>DeepOmix, DeepSigSurvNet, DrugCell, DrugCell+fusion, DrugVNN, DTox, ex-<br>piMap, GCS-Net, GeneticNN, GenNet, GenNet-Interpret, GraphGONet, GSNNs,<br>IRnet, KP-Net, KPNNs, ME+GE, MiNet, MOVIDA, MPAC, MPVNN, Multilevel-GNN, Multitask-VNN, NeST-VNN, OntoVAE, PAAE, PAGE-Net, ParsVNN, PASNet,<br>PathCNN, PathDeep, pathDNN, PathExpSurv, PathME, PBAC, PiDeeL, PINNet,<br>pmVAE, P-Net, P-Net Robustness, priorVAE, ReGeNNe, SigPrimedNet, VEGA,<br>VNNSurv, XAI-AGE, XMR, Varmole |
| No | 7 | BiGMLVQ, DeepBINN, DeepSignalingSynergy, FGNN, GONN, GSAE, k-DNN |

